# Separating three variability and noise sources in the response fluctuation of brain stimulation

**DOI:** 10.1101/2024.11.24.625055

**Authors:** K. Ma, S. M. Goetz

## Abstract

Motor-evoked potentials (MEPs) are among the few directly observable responses to suprathreshold brain stimulation and serve a variety of applications. If the MEP size is graphed over the stimulation strength, they form an input–output (IO), recruitment, or dose–response curve. Previous statistical models with two variability sources inherently consider the small MEPs at the low plateau as part of the neural recruitment properties. However, recent studies demonstrated that small MEP responses are contaminated and over-shadowed by background noise of mostly technical quality and suggested that the recruitment curve should continue below this noise level. This work intends to separate physiological variability from background noise and improve the description of recruitment behaviour. We developed a model with three variability sources and a logarithmic logistic function without a lower plateau. Compared to previous models, we incorporated an additional source for background noise from amplifiers, electrode impedance, and remote bioelectric activity, which form the obesrved low-side plateau. Compared to the dual-variability source modes, our approach better described IO characteristics, evidenced by lower Bayesian Information Criterion scores across all subjects and pulse shapes. The model independently extracted hidden variability information across the stimulated neural system and isolated it from background noise, which led to an accurate estimation of the IO curve parameters. This new model offers a robust tool to analyse brain stimulation IO curves in clinical and experimental neuroscience, reducing the risk of spurious results from inappropriate statistical methods. By providing a more accurate representation of MEP responses and variability sources, this approach advances our understanding of cortical excitability and may improve the assessment of neuromodulation effects.

## I. Introduction

Motor-evoked potentials (MEPs) are among the few directly measurable and observable acute responses to transcranial magnetic stimulation (TMS) of the brain [1], [2]. They therefore serve as a significant biomarker for many applications, such as probing excitability changes through neuromodulation or through the motor threshold as a stimulation reference for almost all other stimulation procedures [1], [3]–[5]. Multiple MEPs administered over a range of stimulation strengths typically form an s-shaped curve with low and high plateaus, also named the motor input–output (IO) curve or recruitment curve, is a well-known and important detection measurement for assessing changes in the motor system [6]–[8]. Peripheral neural and neuromuscular stimulation forms similar IO curves [9]–[12]. IO curves are commonly fit using ordinary least-square regression of sigmoidal curves, such as Boltzmann- and Hill-type functions, which intrinsically assume additive Gaussian errors on top of the MEP size [13]–[16].

However, this simple method is insufficient to model the inherently variable and noisy MEP features. The MEP in response to the brain and certain types of spinal stimulation can exhibit high levels of intra-individual trial-to-trial variability and rather intricate, highly skewed, heteroscedastic distributions [17]–[19] Therefore, the least-square fitting of a linear sigmoid would lead to asymmetric residuals and is prone to spurious results, which could cause systematic errors in the IO curve parameters and derived biomarkers. Instead, logarithmic normalisation of the MEP distribution was strongly recommended and has been suggested to improve stability and reduce model bias when calibration [20], [21].

Beyond normalisation, available statistical models could identify and isolate several widely independent sources of variability involved in IO curves. One additive source on the stimulation side appears to primarily affect the curve in the x direction, i.e., the stimulation-strength direction modulating the neuronally effective stimulus strength. Another variability source is rather independent of the stimulation and multiplicatively changes the MEPs in the y direction, i.e., the MEP-amplitude direction [22]. It can quantitatively describe the size of both variability sources. For example, cortical excitability fluctuations would be exclusively represented by the additive variability source and the spinal and muscular pathways by the multiplicative source [18], [23], [24]. In comparison with the conventional least-square regression model, later research refined the model and confirmed its statistically better description of the observed IO characteristics of changing distribution spread and skewness with stimulus strength [24].

Apart from fluctuations in the corticospinal systems, the background noise also significantly affects the shape of the motor IO curves, which represents the noise emerging in a recording while the electrode is on the skin. It most likely comes from the amplification system, known as thermal noise, and the interface between the skin and the electrode [25], [26]. Additionally, the bioelectric activities from other muscle units that are not targeted might also contribute to the noise floor [27]. For low stimulus strengths, the MEP responses seem to disappear, and the measurement noise dominates the lower plateau of the IO curve [28]. However, a matched-filter signal detector instead of a conventional peak-to-peak reading can decrease the effective noise level and therefore also the lower plateau of the MEP IO curve by as much as an order of magnitude [28], [29]. Since the lower level is primarily determined by technical factors, it should not be included in the IO curve or considered as a biomarker. From another perspective, these studies suggested that, due to the limitations of the amplifier and detection techniques, the background noise conceals weak MEP responses when the neural system actually does respond to the small stimuli. Hence, although capable of fitting and explaining the stimulus-strength-dependent MEP amplitude distributions in previous studies, it appears inappropriate to use an s-shape curve with a lower boundary for simplicity that lumps both the background noise floor and weak MEP responses together.

Therefore, we propose a new statistical model that separates MEP responses and background noise. The model incorporates three variability sources beyond the two previously identified ones: one sits at the input side, modulating the stimulation site or locations where the recruitment information is still available due to interactions of various units, one may act at the corticospinal and muscular pathways, and one acts at the measurement side. Consequently, this model is called a triple-variability-source model. This separation allows us to independently represent the neural properties without the disturbance from the background noise. Furthermore, the variability sources have rather different modes of action (multiplicative vs. additive) and with different distributions. Finally, we decided to remove the sigmoidal recruitment model, specifically its lower plateau, because the lower plateau was found to lack a physiological basis. Instead, it appeared to be an artefact caused by technical issues such as equipment noise and varying electrode impedance during a session. The model can analytically extract hidden information of variability sources over the entire stimulated neural system from the MEP responses. It quantitatively describes the noise floor and identifies the expected MEP amplitudes below the measurement noise floor at lower stimulus strengths.

## II. Methodology

### A. Mathematical framework

Figure 1 depicts the overall structure of this triple-variability-source MEP IO model of the stimulated motor system. The input of the model, *x*, is stimulus strength and its output is MEP amplitude, *y*. The model has two distinct functional blocks. The recruitment block represents how the neural system responds to external stimulation. We used a logarithmic logistic function

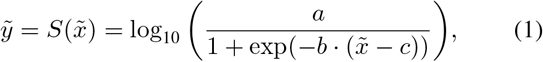

where 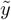 is the output of the recruitment function (see Figure 1), 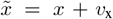 is the effective excitation at the input of the recruitment, and all parameters must be positive values. The exponential transformation block translates the neural responses from the logarithmic domain to the normal scale domain to symmetrise the MEP distribution [3], [20]. The base of the exponential function can be chosen freely because of the logarithmic base change rule. Hence, we used 10 as the base here for simplicity.

**Fig. 1.**
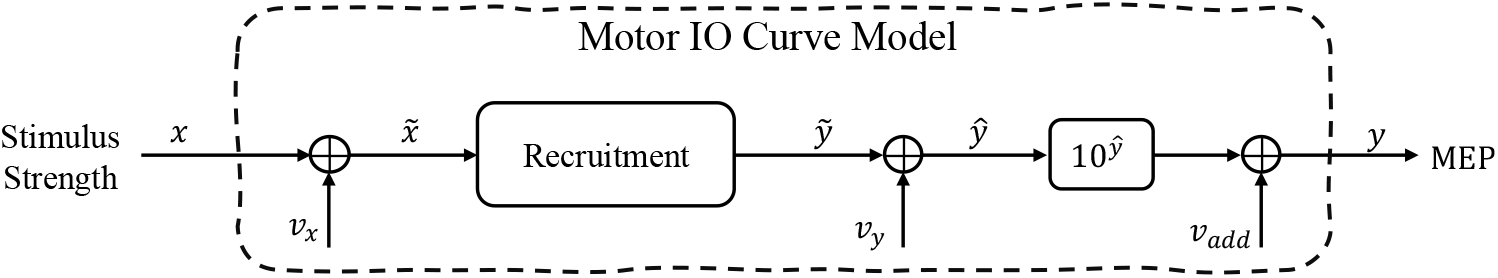
Block diagram of the motor input–output (IO) response model with triple variability sources. Variable *x* is the stimulus strength, recruitment is the neural recruitment characteristics, 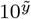 is an exponential transformation block, and *y* is the measured peak-to-peak electromyographic (EMG) amplitude. Independent stochastic variables *v*_x_, *v*_y_, and *v*_add_ with their density functions *g*_x_, *g*_y_, and *g*_add_ represent three different variability sources respectively. 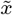 and 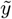 are respectively the input and output of the characteristic function. MEP: motor-evoked potential.

The motor IO curve model has three independent variability sources acting at various locations over the entire neural pathway from the cortex to the muscular motor units. Before the recruitment, the *v*_x_ variability source, as an additive noise, acts along the x axis of the IO curve and can represent the short-term excitability fluctuations of the neurons directly activated by the stimulus. The output-side variability source *v*_y_ before the exponential transformation affects the MEP responses in the logarithmic domain and accordingly has a multiplicative character as shown in Figure 1. This variability source may result from short-term fluctuations in the spinal pathways, the synaptic connection between motor neurons, and the neuromuscular junction and the myocytes [19], [30]. Moreover, recent studies demonstrated that the motor system still can be evoked by weak stimulation and suggested that a minimum response for the cortical neuronal target likely exists but seems far less than the background noise floor [24], [28], [29], [31]. Thus, we introduced an additional variability source along the y axis, *v*_add_, after the exponentiation to represent the additive noise. Therefore, the entire IO curve model becomes

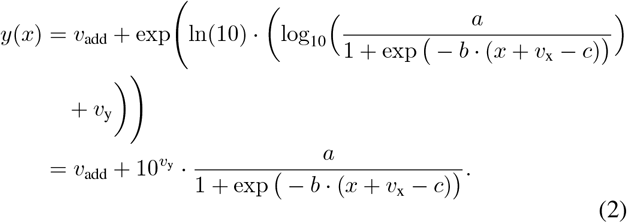

This IO curve models the neural recruitment characteristics with a low-side plateau not as part of recruitment but resulting from the background noise, a monotonically growing section representing the increasing population of activated neurons as the stimulus strength increases, and an upper saturation plateau corresponding to the maximum achievable population responses to the sufficiently strong stimulus. Thus, in comparison with previous recruitment curve models, our curve Equation 1 no longer uses a lower plateau to represent the background noise floor, for which we used another random variable, *v*_add_, instead. Hence, this functional property can independently consider the noise floor and help us easily distinguish it from the small MEP responses in a semi-logarithmic plot.

### B. Model parameters

The triple-variability-source model has two different groups of parameters, one set comes from the logarithmic logistic function and the other set from those three aforementioned variability sources. Fitting this model only requires the measured stimulus–response pairs (*x*_*i*_, *y*_*i*_). Since we need to estimate the parameters of specific probability density distributions, we applied the maximum likelihood estimation to fit the model using such collected data. In the perspective of conditional probability, Equation 2 explains that, for a given *x* and a parameter set of the model ***θ***, the system may output the corresponding *y* with a certain conditional probability, *f*_*y*|*x*_(*y*|*x*, ***θ***). Calculating the corresponding conditional probability requires consideration of the entire IO model. For a fixed stimulus *x*, the input of the recruitment 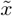 has a probabilistic density function 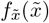, which is equivalent to the density distribution of the variability source 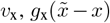. The conditional probability density distribution of the output dependent on 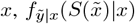, is thus given as

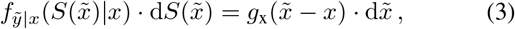

where d· denotes a classical differential.

The second variability source *v*_y_ involved in *ŷ* (see Figure 1) interplays with 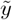. According to the probability convolution for the addition of two independent probability distributions, the density function of *ŷ* given a fixed *x* is

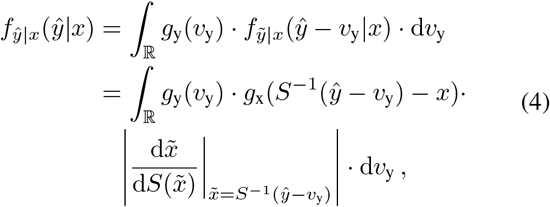

where *S*^−1^(·) is the inverse function of *S*(·) and the value of 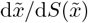 is evaluated at point 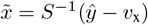.

Likewise, the probability density distribution of 10^*ŷ*^ after exponentiation follows the conversion

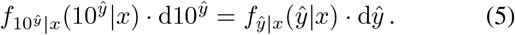

Therefore, the probability density distribution of the output *y* is

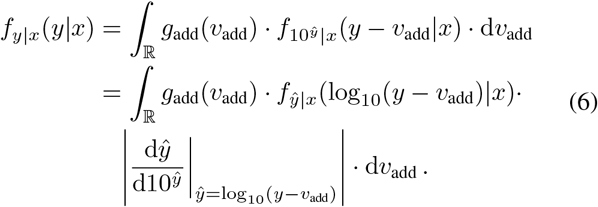

Both variability sources *v*_x_ and *v*_y_ are implemented as independent stochastic elements with Gaussian distribution and respectively obey 𝒩(0, *σ*_x_) and 𝒩 (0, *σ*_y_), where *σ*_x_ and *σ*_y_ are the standard deviations. Previous work suggested that the background noise variability source *v*_add_ transformed by a peak-to-peak extraction algorithm is non-Gaussian but obeys a generalised extreme-value distribution, *GEV* (*k, σ*_add_, *μ*_add_), instead [32]. *k* controls the distribution shape, *σ*_add_ the distribution spread, and *μ*_add_ the distribution location [33]. Accordingly, the entire model parameter vector, which fully describes a subject’s recruitment behaviour to a specific pulse type, is ***θ*** = [*a, b, c, σ*_x_, *σ*_y_, *k, σ*_add_, *μ*_add_].

### C. Data acquisition

We obtained the experimental data from the literature with nineteen healthy subjects (age 18 − 45, 12 female and 7 male, all right-handed) [28], [29]. The data include single-pulse TMS with various pulse shapes, such as monophasic, reverse monophasic, and biphasic. MEPs were recorded and sampled synchronously to the TMS pulse trigger through surface Ag/AgCl electrodes and an MEP amplifier (K800 with SX230FW pre-amplifier, Biometrics Ltd, Gwent, UK) at 5 kHz and 16 bit. The recording starts 200 ms before the delivery of the testing TMS pulse, and the whole duration spans 410 ms.

Before peak-to-peak *V*_pp_ detection, we removed the DC component of the electromyography (EMG) recordings and then filtered the high-frequency components of them with a fourth-order Butterworth filter with a cut-off frequency of 600 Hz. Additionally, recordings with activity of more than 40 μV (peak-to-peak) within a window of 200 ms before the TMS pulse were marked as facilitated and excluded from the analysis [28], [29]. The stimulus strength was expressed as a percentage of the TMS device’s maximum amplitude (% MA).

This study used a self-learning matched-filter algorithm to detect and extract *V*_pp_ with increased sensitivity [29]. We set the sliding filter detection window to 20 ms. By visually checking the active MEP responses, the detection window runs from 220 ms to 260 ms, decomposes the active responses into the summation of several motor unit action potentials, and finds the peak-to-peak voltage (see the literature for details [29]). Moreover, we applied the same filter to measure the background noise by going through the first 40 ms of each MEP measurement.

### D. Parameter calibration

Each subject *i* has a set of *N*_*ij*_ stimulus–response pairs {(*x*_*ijk*_, *V*_pp,*ijk*_) | *k* ∈ [1, *N*_*ij*_], *k* ∈ ℤ^+^} for each pulse shape *j*. A specific subject shares the same parameters for *v*_add_ across different pulse shapes, and the parameters of *v*_add_ are firstly estimated by using the likelihood function of GEV distribution based on the measured background noise. Since the *V*_pp_ detection algorithm must generate positive amplitude values for MEP measurements, that is, the GEV distribution should always be positive. We set up a constraint that forces the distribution to equal zero when the random variable is less than zero. According to Equation 6, the remaining parameters of ***θ*** (by keeping the parameters of GEV constant) for each subject and each pulse shape are estimated through

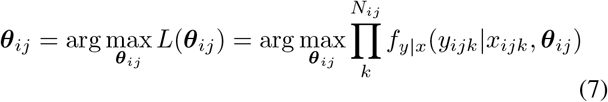

under the assumption that the inter-pulse correlation with sufficient temporal spacing is negligible [13]. The likelihood maximisation for the objective function 7 is achieved with a global particle-swarm optimisation algorithm, and all codes are implemented using MATLAB (USA, v2022b). All codes are available online (Supplementary material).

### E. Model validation, comparison and analysis

Before calibration on the experimental dataset, we tested the performance of the particle swarm optimisation algorithm. We simulated the process based on Equation 2 with the theoretical parameters ***θ*** = [1, 0.45, 60, 3, 0.2, 0.1, 1 × 10^−3^, 3 × 10^−3^] and generated 500 data points with 25 different stimulus strengths spanning from 30% to 100% of the device’s maximum amplitude. There are 20 measurements for each given stimulus strength *x*. In addition, we used a GEV random number generator to generate 1, 000 background noise data points. To validate the robustness of the optimisation performance and the reliability of model accuracy, we implemented a K-fold cross-validation approach with *K* = 30. In each iteration, 29 subsets were used to calibrate the model. This method assessed the model’s generalisability and provided reliable estimates of its performance and parameter stability.

We employed mixed-effect models to analyse the calibrated model parameters, accounting for biological differences among individuals and variations in experimental techniques. For parameters [*a, b, c, σ*_x_, *σ*_y_], we selected *age* (continuous), *sex* (categorical), and *pulse shape* (categorical) as fixed-effect variables, with *subject* as a random intercept variable. For parameters [*k, σ*_add_, *μ*_add_], we used a different approach. Since pulse shape does not affect background noise, we excluded it from these models. Additionally, the number of subjects equalled the number of sets of GEV parameters, making it insufficient to serve as a random effect. Consequently, we employed simple linear models to analyse these GEV parameters. We used the *lme4* package in R (version 4.3.1) to calibrate the mixed-effect models [34]. To test the significance of the fixed-effect variables, we conducted a type-III ANOVA with Satterwaite’s method. Moreover, we used the likelihood-ratio test to determine the significance of *subject* for parameters. In total, we developed and analysed eight distinct models: five mixed-effect models for [*a, b, c, σ*_x_, *σ*_y_] and three linear models for [*k, σ*_add_, *μ*_add_].

Moreover, a previously proposed dual-variability-source model from a decade ago has two variability sources at the input and output sides to fit the MEP IO curve [22]. We calibrated this dual-variability-source model for each subject as well as each pulse shape and used the Bayesian information criterion (BIC) as an indicator of goodness-of-fit to support model comparison [35]. This comparison is appropriate because the BIC takes into account not only the number of parameters of the statistical model but also the quantity of the calibration data. Moreover, we resampled the estimated model parameters and their BIC values using boot-strapping analysis with 1, 000 repetitions for deriving their median and interquartile range. For statistical significance testing, we employed the Mann-Whitney U test for single comparisons and post-hoc analysis. We set significance levels at *α*_double_ = 0.05 for double-sided tests and *α*_single_ = 0.025 for single-sided tests.

## III. Results

### A. Model validation

To clearly present the physiological properties and technical noise of IO curves, we separately plotted the recruitment curves and their corresponding noise floor. Figure 2 demonstrates a representative triple-variability-source model calibrated to the simulated dataset. Two variability ranges, i.e., 85% and 95%, represent the overall IO trend and encompass most of the simulated data points. The expected neuronal recruitment curve (solid red line) closely follows the trend of the simulated data points and captures the shape of the input–output relationship in responses. The background noise level, represented by the mode value of the variability source *v*_add_ (dashed red line), is clearly distinguishable from the recruitment curve. The mode value well represents the background noise because the sample points are denser around the mode value over the entire variable domain.

**Fig. 2.**
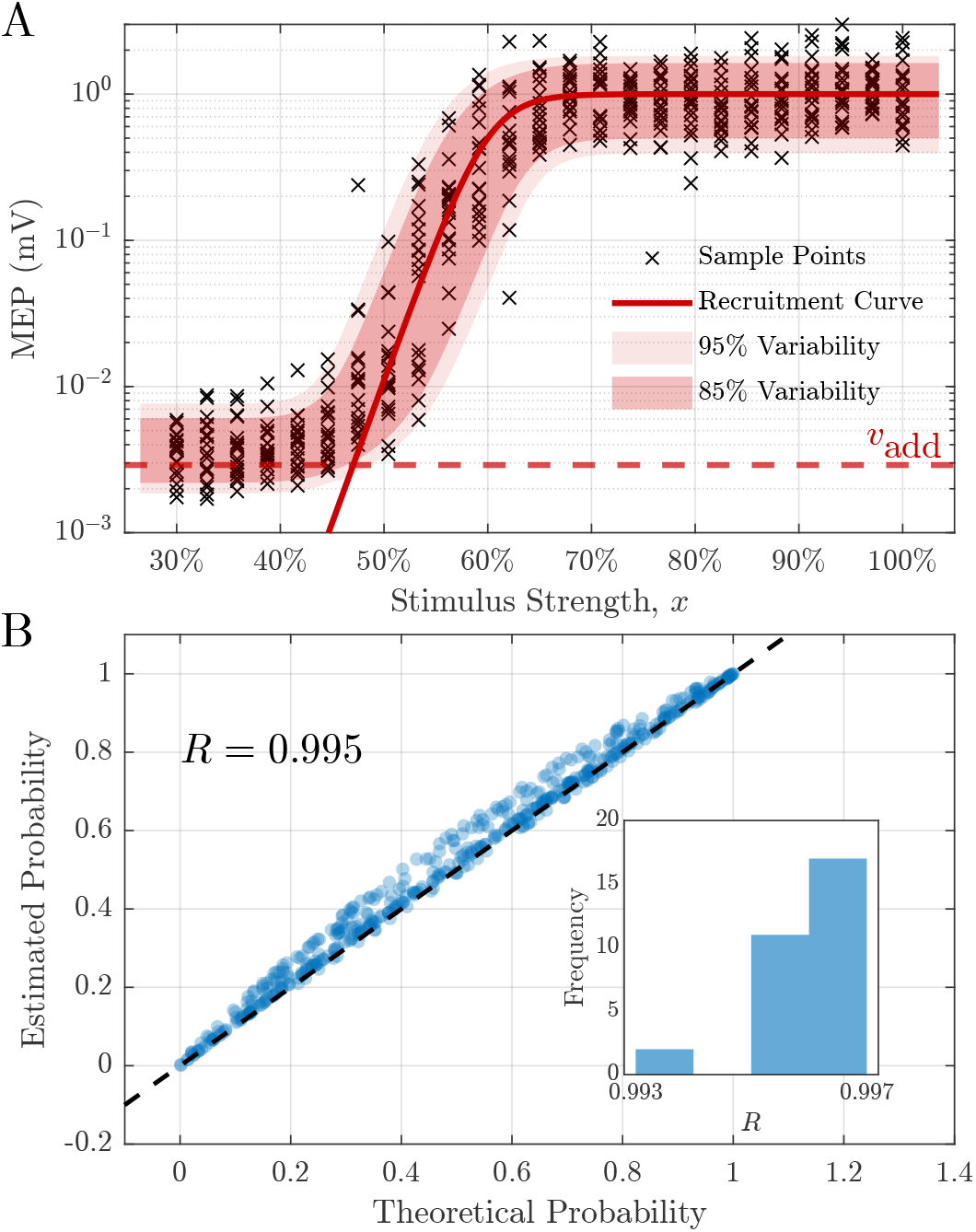
Representative triple-variability-source model calibrated to simulated data with known properties. (**A**) Calibrated curve model consisting of a background noise level (red dashed line, *v*_add_) and an expected neuronal recruitment curve (red solid line). The two variability ranges of Equation 2 are 95% (shaded with light red) and 85% (shaded with moderate red). (**B**) Comparison of the cumulative probability between the theoretical models (theoretical probability) and the representative estimated model (estimated probability) with a correlation coefficient of *R* = 0.995. The inset represents the distribution of correlation coefficients between the theoretical and estimated probability for *K* = 30 cross-validated models. The parameters used in this figure are *a* = 0.98, *b* = 0.46, *c* = 60.50, *σ*_x_ = 2.93, *σ*_y_ = 0.20, *k* = 1.07 × 10^−1^, *μ*_add_ = 1.01 × 10^−3^, and *σ*_add_ = 3.04 × 10^−3^.

According to Equation 1, the neuronal recruitment curve almost equals 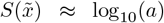 when 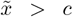, whereas 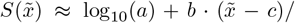 ln 10 when 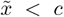. Thus, *c* acts as a junction and divides the neuronal recruitment curve into transition and saturation regions. The curve in the transition region rapidly increases over a small range of stimulus strengths with a slope *b* from the noise level and becomes saturated at a level of *a* in the saturation region. Therefore, we can respectively define [*a, b, c*] as the saturation level, slope, and shift of the neuronal recruitment curve.

Table I lists the estimated parameters with their median and interquartile values. The estimated parameters closely match their theoretical value with an average relative deviation on the order of 10^−2^ or lower and have narrow interquartile ranges as low as 10^−3^, which indicates high accuracy and precision in the estimates. Figure 2 (**B**) contains a probability–probability (P–P) plot with the estimated and theoretical parameters to calculate the corresponding cumulative probabilities of the triple-variability-source model and illustrates that most points align with the diagonal dashed line (*R* = 0.995), which suggests an excellent agreement between the theoretical and the estimated model. In addition, the distribution of the correlation coefficient (*R*) of the P–P plots for these cross-validated models spans a very narrow range from 0.993 to 0.997 (inset of Figure 2 (**B**)). These results suggest that the optimisation procedure successfully recovered the underlying parameters used to generate the simulated data and validates the optimisation robustness, intrinsic model reliability, and reproducibility with respect to deterministic curve parameters and variability descriptors.

**TABLE 1.** Comparison of Theoretical and Estimated Model Parameters.

| | $a$ | $b$ | $c$ | $\sigma_x$ | $\sigma_y$ |
| --- | --- | --- | --- | --- | --- |
| Theoretical value | 1 | 0.45 | 60 | 3 | 0.2 |
| Estimated value | 0.97<br>($1.07 \times 10^{-2}$) | 0.47<br>($7.64 \times 10^{-3}$) | 60.32<br>($1.48 \times 10^{-1}$) | 2.96<br>($4.70 \times 10^{-2}$) | 0.20<br>($2.61 \times 10^{-3}$) |

Note: Estimated values are given as median and interquartile range (in the parentheses). The GEV parameters were estimated using simulated baseline background noise and kept the same in each iteration across the cross-validation; they were $k = 1.07 \times 10^{-1}$ , $\mu_{add} = 1.01 \times 10^{-3}$ , and $\sigma_{add} = 3.04 \times 10^{-3}$ .

### B. Model regression results

Figure 3 graphs a calibrated model for Subject S15A as a representative example; the data for all subjects is available in the Supplementary material. Figure 3 (**A**) demonstrates that the neuronal recruitment curve describes the overall trend of the strength-dependent responses, and its two variability ranges encompass the majority of the sample points. Figure 3 (**B**) shows the distributions of *v*_x_ and *v*_y_ for different pulse shapes, and Figure 3 (**C**) illustrates that the distribution of the background noise, *v*_add_, which forms the low-side plateau, is highly right-skewed and has a long tail towards larger response values. The calibrated GEV model closely matches the distribution of the measurements. In addition, it is an exclusively positive distribution and vanishes below zero. Thus, noise never reduces the MEP amplitude reading in a peak-to-peak detector. This behaviour reflects practical observations.

**Fig. 3.**
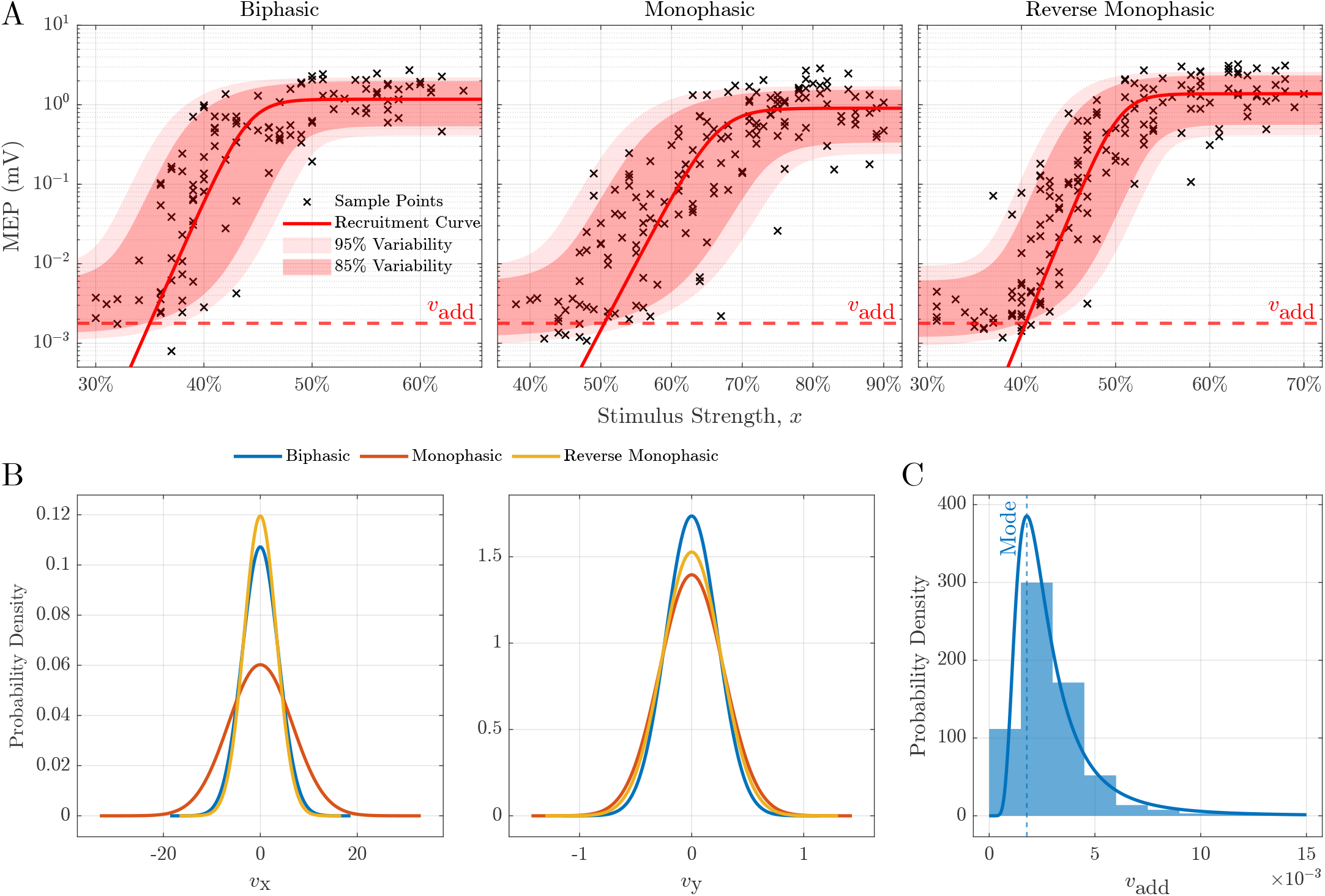
Representative individual IO data, regression results, and distributions of their variability sources for Subject S15A. (**A**) Illustration of the two components of the triple-variability-source model for each pulse shape, an expected neuronal recruitment curve (red solid line) and a mode value of the background noise *v*_add_ (red dashed line); the x axis represents the TMS pulse strength (percentage of the maximum amplitude of the device). The y axis is the peak-to-peak amplitude of the MEPs. The measured MEP amplitudes to a stimulus are marked with crosses (×). The two variability ranges of Equation 2 are 95% (shaded with light red) and 85% (shaded with moderate red). (**B**) Distribution of variability sources *v*_x_ and *v*_y_. (**C**) Distribution of the GEV model of *v*_add_ and the corresponding baseline measurements for the subject; the vertical blue dashed line is the mode value of the GEV model.

Moreover, correlation analysis demonstrated a moderate negative correlation between parameters *b* and *c* (*R* = −0.43, *p* = 8.94 × 10^−4^), as they are associated and may partly compensate each other in Equation 1. This correlation suggested that an increased value of the motor threshold may decrease the value of the ascending slope. In addition, these recruitment curve parameters are also related to variability sources. The strongest (negative) correlation was observed between parameter *b* and variability *σ*_x_ (*R* = −0.77, *p* = 2.67 × 10^−12^). In contrast, parameter *c* showed a strong positive correlation with *σ*_x_ (*R* = 0.60, *p* = 8.78 × 10^−7^). These opposite relationships are further supported by the correlation between *b* and *c*. Additionally, the saturation level *a* demonstrated a strong negative correlation with variability *σ*_y_ (*R* = −0.73, *p* = 1.56 × 10^−10^), indicating that higher saturation levels are associated with more consistent muscle unit responses.

Previous research suggested that the curve slope may be a by-product of different thresholds rather than a viable biomarker [7], which aligns with our observed correlation between slope *b* and shift *c*. The correlation between *b* and *σ*_x_ appears to be a consequence of this relationship, suggesting that both slope *b* and variability *v*_x_ are secondary effects of parameter *c*. Likewise, variability *v*_y_ would be a secondary effect of parameter *a*. To account for these dependencies and avoid potential misinterpretation, we normalised both *b* and *σ*_x_ by *c* and *σ*_y_ by *a* before conducting significance analysis.

Mixed-effects analysis demonstrates that the factor *pulse shape* significantly influences model parameters, while neither *age* nor *sex* show significant impact on any parameter. Table II summarises the variability sources and recruitment parameters for different pulse shapes derived with boot-strapping. The MEP saturation level varies moderately between pulse shapes (*p* = 0.04, *F* (2, 35.05) = 3.53). Biphasic pulses generate the largest saturation level, comparable to reverse monophasic pulses (*U* = 168, *p* = 0.94). Monophasic pulses with normal current direction demonstrate smaller saturation levels compared to reverse monophasic pulses (*U*_left_ = 118, *p* = 0.03). Whereas normalisation eliminates differences in the slope 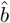 (*p* = 0.15, *F* (2, 35.56) = 1.97), the pulse shape significantly shifts the IO curves (*p* = 1.87 × 10^−15^, *F* (2, 35.13) = 103.52). Post-hoc analysis confirms that monophasic pulses with normal current direction shift the curve rightward compared to both biphasic (*U*_right_ = 516, *p* = 1.3 × 10^−6^) and reverse monophasic pulses (*U*_right_ = 517, *p* = 1.0 × 10^−5^), reflecting known threshold differences between pulse shapes.

**TABLE 2.**
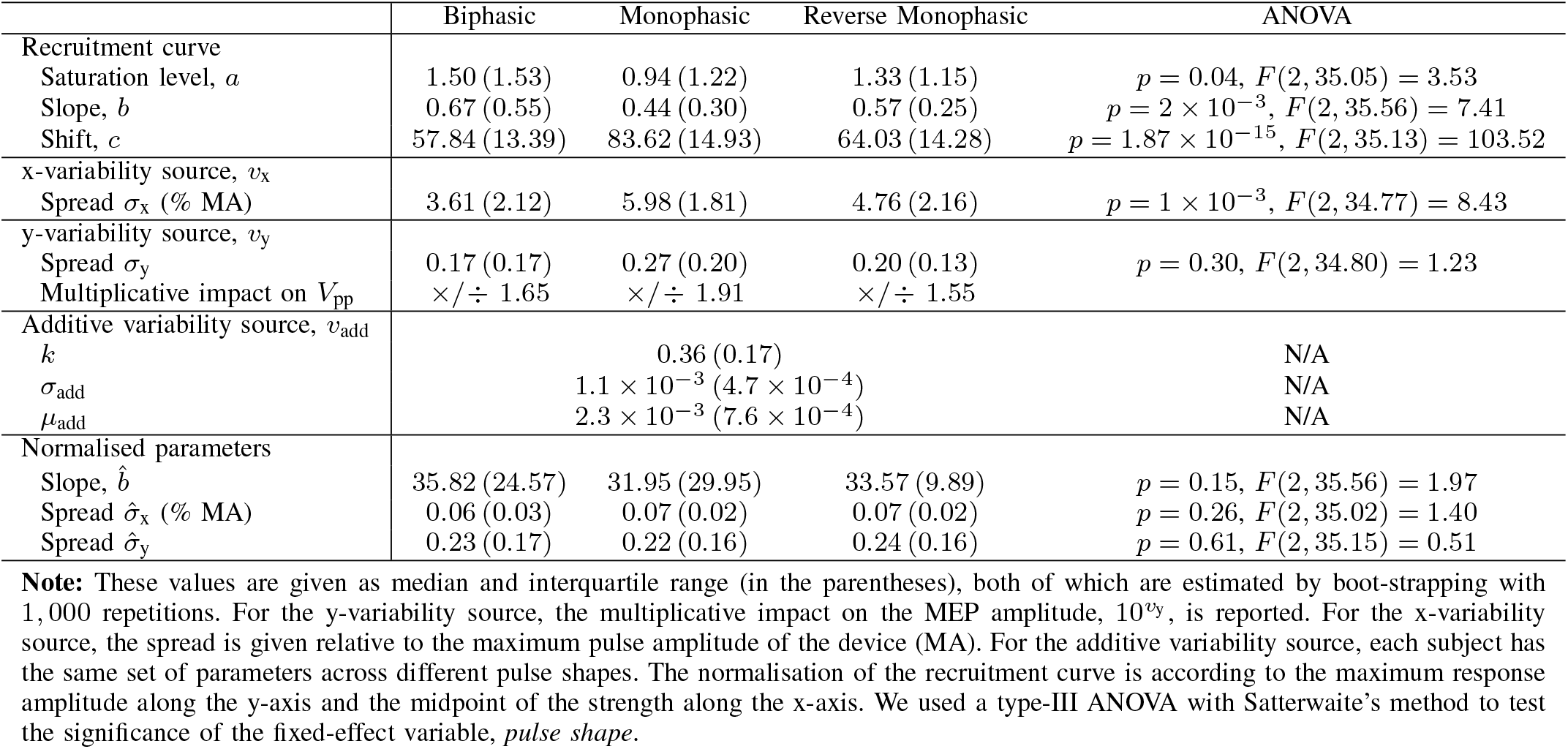
Summary of the estimated IO curve variability and recruitment parameters for the triple-variability-source model.

The x-variability source, expressed as a percentage of the maximum TMS device output strength, shows no differences between pulse shapes after normalisation (*p* = 0.26, *F* (2, 35.02) = 1.40). Similarly, the y-variability shows no significant differences for either original (*p* = 0.30, *F* (2, 34.80) = 1.23) or normalised values (*p* = 0.61, *F* (2, 35.15) = 0.51). The additive variability source, *v*_add_, exhibits a right-skewed distribution (*k >* 0) across all subjects, with values concentrated on the left side. With *μ*_add_ = 2.3 × 10^−3^ and *σ*_add_ = 1.1 × 10^−3^, the background noise primarily occurs at low magnitudes of 10^−3^ mV within a narrow range, potentially obscuring MEPs with responses at or below this noise level.

Finally, likelihood-ratio tests reveal significant subject-specific effects on parameters *a* (*χ*^2^ = 23.47, *p* = 1.27 ×10^−6^), *c* (*χ*^2^ = 26.85, *p* = 2.19 × 10^−7^), and 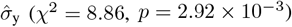. These results suggest that saturation level, curve shift, and variability *v*_y_ are likely individual features that explain inter-subject variability.

### C. Comparison with the dual-variability-source model

Figure 4 (**A**) presents three representative regression results for both the dual-variability-source and the triple-variability-source model of three subjects for a visual comparison between them. First, the dual-variability-source model consistently estimates the saturation levels rather poorly compared to the triple-variability-source model. For instance, in the case of the reverse monophasic pulse of S08A (Figure 4 (**A**), left panel), the tripe-variability-source model saturates for every stimulus strength over *x ≈* 60 %, whereas the dual-variability-source curve still increases after *x ≈* 80 %. The dual-variability model may also be struggling with poor convergence considering its higher number of degrees of freedom in the sigmoid description.

**Fig. 4.**
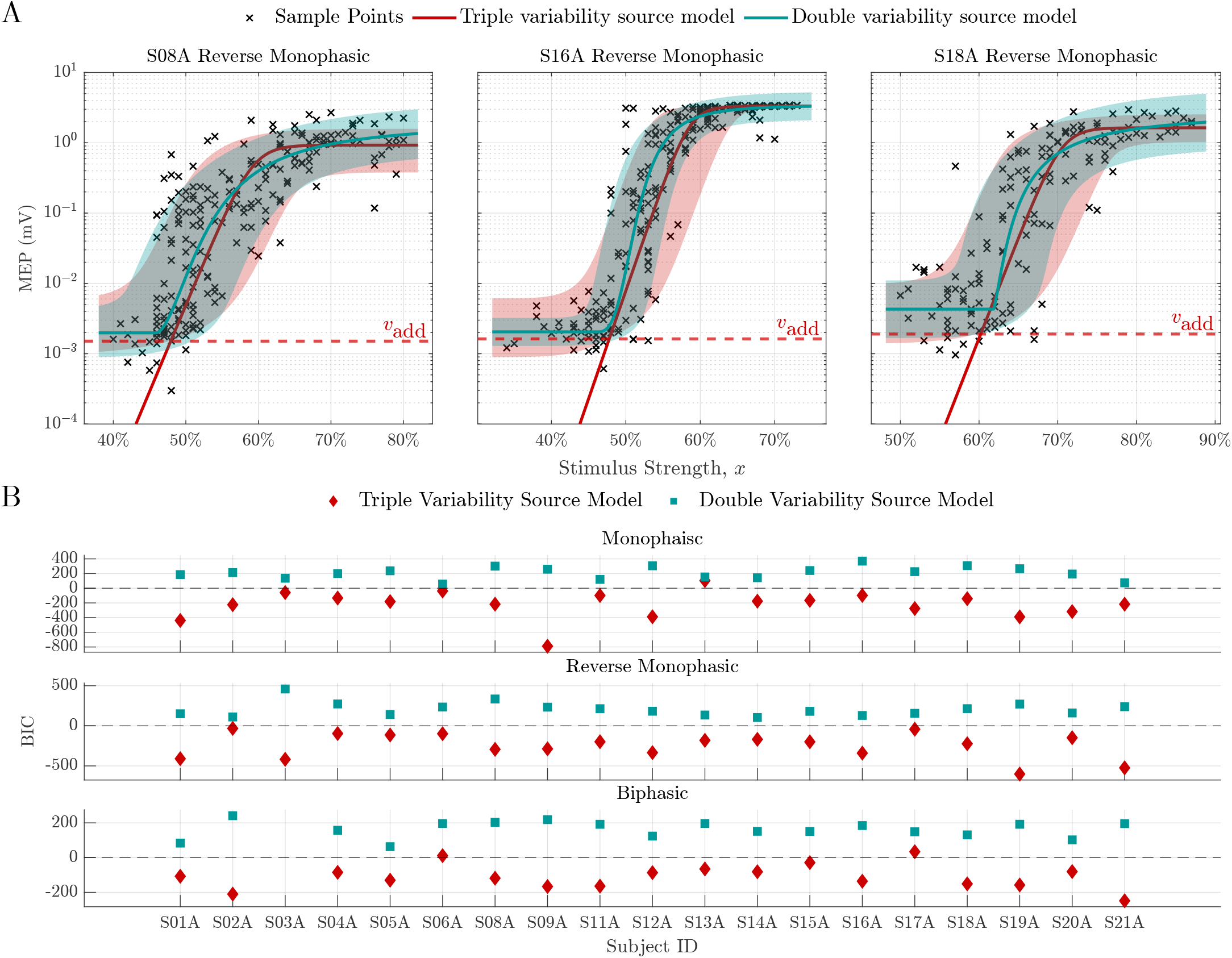
Representative individual IO data and regression results for dual- and triple-variability-source models. (**A**) Regression results from three selected subjects for both models (red: triple-variability-source model; teal: dual-variability-source model). The x-axis represents the TMS pulse strength as a percentage of the maximum amplitude of the device. The y-axis is the peak-to-peak amplitude of the MEPs. The measured MEP amplitudes in response to stimuli are marked with crosses (×). Each neuronal recruitment curve is associated with a 85% variability range shaded with red (triple-variability-source) or teal (dual-variability-source) colour. (**B**) Bayesian information criterion (BIC) for both models separately (red diamond: triple-variability-source model; teal square: dual-variability-source model).

Moreover, the dual-variability-source model inherently integrates both y-variability sources, specifically the MEP fluctuations and the additive recording noise along the y axis, in the same *σ*_y_, although they differ substantially in size and shape. Thus, the regression results even depend on sampling. For example, regarding the case of S16A (Figure 4 (**A**), middle panel), the dual-variability-source model estimates a narrow variability range for the noise level because there are more concentrated measurements gathering at the saturation level. Likewise, a higher number of sparse measurements at the noise level likely causes the dual-variability-source model to overestimate the variability range of the saturation level in the case of S18A (Figure 4 (**A**), right panel). Since background noise and variability as two different effects are lumped together, the sampling, i.e., the number of pulses close to the noise floor vs. of pulses at high stimulation strength determines the estimated variability. In contrast, the triple-variability-source model introduces an additional source *v*_add_ that specifically accounts for device and physiological noise. Thus, the new model can independently estimate the variability ranges of the saturation and noise levels according to the measurements and precisely capture the distribution differences, as demonstrated in all cases in Figure 4 (**A**).

Another notable difference is observed in the varying rise slope of the dual-variability-source model, whereas the triplevariability-source model increases with a less steep constant rate in logarithmic scale from the background noise floor before reaching the saturation level. However, in some cases, e.g., S18A (Figure 4 (**A**), right panel), the motor system responds to the TMS stimulation before the rise of the dual-variability-source model, a scenario well captured by the triple-variability-source model. Additionally, regarding the estimation of background noise, the triple-variability-source model consistently yields lower levels than the dual-variability-source model, with more sample points clustering around the mode value of *v*_add_ rather than the low-side plateau in the dual-variability-source model.

Table III summarises only the variability sources across all pulse shapes and provides the BIC scores for both the dual-variability-source and triple-variability-source models for comparing the goodness-of-fit. The dual-variability-source model estimates variability values similar to those of the triple-variability-source model. However, the regression of IO data with the triple-variability-source model achieves a significantly lower BIC score than the dual-variability-source model (*U* = 1, 602, *p* = 1.0 × 10^−19^), which demonstrates a higher descriptive quality without over-fitting. This result suggests that the logarithmic logistic recruitment curve outperforms the original sigmoidal function and better describes the trend of MEP IO curves.

**TABLE 3.**
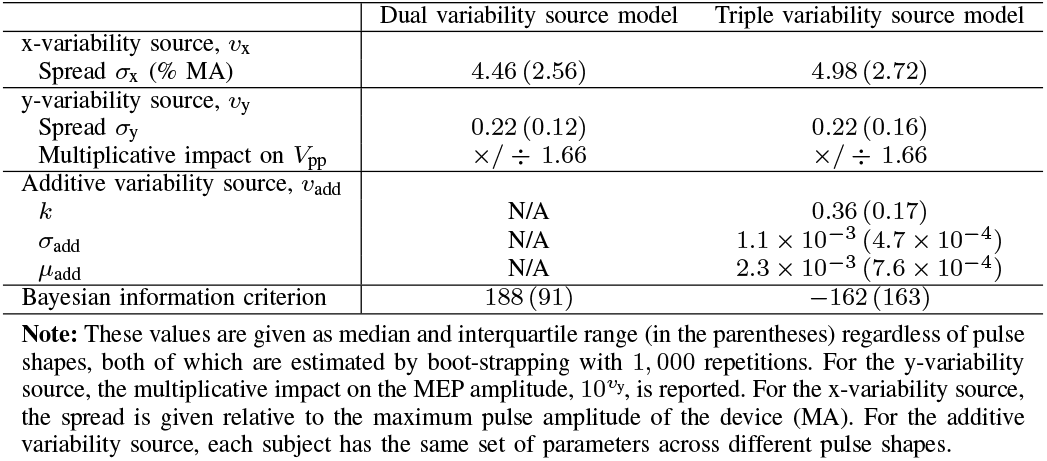
Comparison of the estimated IO curve variability and goodness-of-fit for the dual-variability-source model and the triple-variability-source model.

## IV. Discussion

Accurate statistical models that can be trained to experimental MEP data and represent the variability accurately can help explain the physiology, extract individual motor system properties, and separate the variability into its components. The variability is not necessarily noise or random. Endogenous signals received by the neuronal target also change the excitability for a short duration and can shift a stimulus–response pair in the IO graph.

### A. Comparison with previous models

Compared with the previous dual-variability-source model, this study used a different recruitment curve without a lower plateau. The lower plateau of the previously typically sigmoidal function was found to not represent a physiological property but rather a technical one, specifically the recording noise floor. Accordingly, this technical property, which would change for an individual based on equipment and set-up, previously led to a different set of IO curve parameters. We therefore separated the low-side plateau and introduced an additional variability as an additive source to represent the background noise. This model could describe varying MEP distributions across the range of stimulus strengths and demonstrate significant improvements over all previous models, including those with already two variability sources.

### B. Physiological interpretation of variability sources

The various variability sources are separated by their different properties and different interactions with the stimulation strength. The different dependence on the stimulation strength indicates different mechanisms and potential locations in the motor system. The variability acting on the input side of the recruitment model with an exclusive effect in the stimulus-strength direction (i.e., along the x axis) represents fluctuations of the effective stimulation strength. Such interaction can only occur at the site where the electric field of the TMS coil activates neurons, which then individually respond with an all or none signal, or at places where the signals of various motor units may interact and recruitment information is still available, e.g., the spine, where several motor units can interact synaptically.

Random noise at the membrane of stimulated neurons [30], [36] would be subsumed in this variability term just as any coil position and orientation fluctuations, which would lead to changing electric fields at the stimulation target from stimulus to stimulus [37]–[40]. However, these effects are known to not explain the observed variability alone: neither more consistency of coil positioning nor invasive electrical stimulation with spatially fixed electrodes could eliminate the pulse-to-pulse variability of MEP amplitudes [23], [41]. This persistent endogenous variability indicates that the x variability of MEPs might be dominated by neurophysiological factors such as undulant endogenous excitability changes in the motor system.

The IO data shows that the multiplicative y variability cannot be influenced by increasing the stimulus strength and still causes variance along the entire IO curve, including the high-side plateau (Figure 3). This result suggests that this variability is independent of the stimulation strength or recruitment and would not be governed by the cortex excitability [18]. Thus, the multiplicative y variability source might be attributed to fluctuations in the motor pathway, e.g., from the spinal cord to and in the muscle cells, where the individual units are supposed to have little interaction in healthy individuals [19]. Moreover, this particular stimulation-strength-independent y is independent of the recruitment, highly skewed but widely normalises under logarithmic transformation, and affects the MEPs all the way to the upper saturation level. Thus, a multiplicative variability term with a simple Gaussian distribution represents this effect well and may indicate that it does not add to but rather modulates an existing signal of an individual motor unit after recruitment in a neuron, a synaptic transmission, or muscle cell. The y variability should further also include latency fluctuations of signals in the individual motor units from the spinal cord onwards, where they do not interact with each other anymore.

### C. Advantages of the triple-variability-source model

In contrast to the dual-variability-source model, the triple-variability-source model introduces a logarithmic logistic function as the recruitment curve and incorporates an additional, additive variability source after the exponential transformation at the model output (Figure 1). This logarithmic function does not have a lower plateau but continues to fall, which reflects the observation of recent studies that MEP responses appear well below this level but are contaminated and over-shadowed by background noise [28], [29]. As in many surface electromyography measurements, the background noise is inevitable and adds independently to the output variability [42], [43]. This noise dominates for small MEPs and masks responses below a certain level [28], [29]. Benefiting from the absence of a lower plateau, this new recruitment curve provides a mathematical basis for incorporating an additional independent noise floor, allowing it to capture distribution differences between the higher and lower sides (for example, S16A shown in Figure 4). Table III demonstrates a better performance of the logarithmic logistic function compared to the original sigmoidal function.

Moreover, this noise is accordingly not a physiological aspect but was accounted for in the multiplicative y variability in previous models. The dual-variability-source model describes both the background noise and the fluctuations in the motor pathways, which therefore is a mix of physiology as well as technology and furthermore depends on relatively similar sampling in the low-side and the high-side plateaus [24]. The resulting variance is a mix of both and does neither correctly represent the noise nor the physiological y variability. Furthermore, the background noise is not only additive but also has a different, very characteristic distribution due to the peak-to-peak MEP detection. Thus, the triple-variability-source model allows for a more accurate and independent estimation of the multiplicative variability inherent in the motor systems as well as the background noise from the perspective of principles of mathematics.

As a result of the combination of various variability effects that share little similarity in one and by wrongly assuming a physiologic low-side plateau, although the IO curve was found to continue falling below that level, the dual-variability-source model lacks robustness when calibrated to data. When the MEP measurements are sparse at the lower stimulus strengths, the dual-variability-source model can entail convergence problems and run into local minima [44]. The triple-variability-source model can instead exploit the statistical and causal independence of the background noise for a statistically more stable sequential approach to estimate it and its entire distribution (i.e., the GEV with all parameters) beforehand. Subsequently, it calibrates all remaining model parameters. We demonstrated this approach here.

In addition to the higher robustness, the background noise estimation can use readily available additional data: measuring background noise is relatively easy compared to MEP response measurement. It does not even need TMS pulses administered and can be extracted from as little as several seconds of background recordings before or between stimulation. Alternatively, the triple-variability-source model can also estimate the entire parameter space at once if the MEP IO database is well-recorded and has enough data points along the entire IO curve, i.e., from just background noise all the way into the saturation level.

### D. Implications for IO curve biomarkers

Many studies on MEPs did not systematically consider the entire range of stimulus strengths together but instead calculated their statistics for individual stimulus strengths and performed pairwise statistical tests or similar group analyses [8], [45], [46]. However, conventional methods may result in spurious results and wrong data interpretation due to unequal and skewed variabilities [18]. The slope, previously reported as a potential biomarker for various aspects, may be an example. A normalisation of the recruitment curve to the parameter *c*, which is similar to converting stimulation strength relative to the motor threshold, eliminated all apparent slope differences (see Table II). The risk that the absolute slope value may be misinterpreted as an independent biomarker but in many cases might rather reflect threshold differences was observed and reported before in several studies [7], [24]. Normalisation by either the motor threshold or a comparable point on the IO curve eliminated all detectable differences in the slope. Accordingly, the slope may to a large share be a secondary impact of other parameters, i.e., likely a threshold effect, and may therefore not be the best and most robust biomarker for physiological analysis.

## V. Conclusion

We proposed a new MEP IO model that separates the rather different trial-to-trial variability phenomena along the motor pathway from the cortex through the spine to the muscle into three widely independent sources and provided a framework to extract the parameters from measurement data for an individual subject. Compared to previous models, the triple-variability-source model could more accurately estimate the MEP IO curve and is less prone to bias or over-fitting. Furthermore, as it reliably extracts quantitative information not only about the expected recruitment but also the variability sources, it offers a tool to study the physiology of MEPs, excitation, and neuromodulation. Furthermore, it can serve as a scientific tool to analyse brain stimulation IO curves in clinical and experimental neuroscience without the previous spurious results and statistical bias.

## Author contribution

KM and SMG envisioned and designed the overall project. KM wrote the code and analysed the data. SL and MQ analysed the data. SMG acquired the data and supervised this project. KM and SMG wrote and edited the text. SL and MQ edited the text.

## Supplementary material

Supplementary documents to this article can be found online (https://github.com/BIOMAKE/Triple Variability Source IO Model).

